# Trait dissimilarity-based tree species loss affects tree diversity effects on herbivory

**DOI:** 10.64898/2026.05.21.726831

**Authors:** Mareike Mittag, Georg Albert, Pablo Castro Sánchez-Bermejo, Andréa Davrinche, Sylvia Haider, Shan Li, Xiaojuan Liu, Ming-Qiang Wang, Andreas Schuldt, Jana S. Petermann

## Abstract

Biodiversity loss can alter interactions not only through changes in tree species richness, but also through the loss of particular functional strategies from ecological communities. Working in a subtropical forest diversity experiment we asked whether tree species richness effects on arthropod herbivory and leaf pathogen infestation depend on community functional diversity, and whether trait dissimilarity-based, non-random species loss alters these relationships compared to random loss. To address this, we combined already established planted scenarios with newly constructed extinction pathways.
We tested the responses of herbivory and leaf pathogen infestation (i) to tree species richness, functional diversity (Rao’s Q), community structure and resource strategies (i.e. evergreenness) and community-weighted trait means as well as predation, and (ii) trait dissimilarity-based extinction pathway analyses that contrasted directed loss of functionally similar versus functionally distinct tree species.
Herbivory increased with tree species richness and this increase was significantly stronger in communities with higher tree functional diversity. Under directed species loss scenarios, herbivory differed most strongly from random-loss expectations when similar tree species were lost first. By contrast, losing functionally distinct species first produced richness effects that were much closer to the random-loss scenarios. Trait-based species loss will therefore modify trophic interactions more strongly than random loss. For pathogen infestation tree richness effects depended on evergreenness and among planted extinction scenarios (three-way interaction), with only minor deviations of trait-based extinction pathways from random-loss expectations. Pathogen infestation also tended to increase with community-weighted mean leaf nitrogen. Predation showed no clear relationship with tree species richness or functional diversity but was positively associated with herbivory. The strength of this association differed among extinction scenarios, providing no evidence for consistent top-down regulation.

**Synthesis:** The ecological consequences of biodiversity loss for leaf damage depend on which functional strategies are lost, not only on how many tree species remain. By integrating observed tree diversity gradients with trait-based extinction pathways, this study shows that functional diversity and host redundancy help explain why herbivores and pathogens are shaped by the same changes in tree diversity through different functional constraints and improve predictions of interaction strength under non-random species loss.

## INTRODUCTION

Biological diversity is declining rapidly worldwide, and this loss has far-reaching consequences for the functioning of ecosystems (Isbell et al. 2011; Hooper et al. 2012; Cowie et al. 2022). As species go extinct, not only functional diversity changes, but also trophic interactions are re-shaped (Dunne et al. 2002; Dirzo et al. 2014). The loss of taxonomic and functional diversity can thus have major consequences, for example by changing interactions between plants and their consumers that directly regulate plant damage and growth performance (Grossman et al. 2018). Forests in particular have been strongly affected by pest species in recent years (Seidl et al. 2017), underlining the importance of understanding how biodiversity loss contributes to interactions between trees and their pests and pathogens. In fact, the diversity of tree species may drive the feeding behavior of herbivores and pathogens and their interactions with trees (Schuldt et al. 2017; Jactel et al. 2021; Rutten et al. 2021; Wan et al. 2026), and ultimately the activity and top-down effects of higher trophic levels, such as predators and parasitoids (Langellotto & Denno 2004; Schuldt et al. 2019). Importantly, changes in tree community composition can also alter herbivory and related functions not only via tree species richness, but also via shifts in functional traits (Schuldt et al. 2014; Wang et al. 2020; Michalko et al. 2024). By reducing functional trait diversity, biodiversity loss can restructure bottom-up and top-down controls and thereby may increase the likelihood of substantial herbivore or pathogen damage in tree-species-poor communities (Seidl et al. 2017; Grünig et al. 2020).

Tree diversity experiments are particularly useful in identifying the consequences of biodiversity loss and have shown that insect herbivory and pathogen damage often change systematically with tree species richness, but the direction and strength of tree species richness effects vary among studies and systems (Schuldt et al. 2015; Grossman et al. 2019; Jactel et al. 2021). This variation is increasingly attributed to community composition and associated leaf traits that shape host apparency, resource concentration, leaf quality and defense, as well as top-down control (Coley & Barone 1996; Wright et al. 2004; Wang et al. 2020). Functional traits hence establish an important link between tree diversity and the bottom-up and top-down regulation of interactions between plants, herbivores, and predators. In forests dominated by specialist herbivores, herbivory often declines with increasing tree species richness, consistent with reduced host concentration and leaf trait-mediated decreases in host suitability (Jactel & Brockerhoff 2007), whereas in forests dominated by generalists, herbivory can remain unchanged or increase, presumably because mixed stands provide a broader diet and complementary resources (Jactel & Brockerhoff 2007; Wein et al. 2016). Evidence from a tree diversity experiment in subtropical China (BEF China) suggests that herbivore community responses can be shaped by a strong contribution of generalist herbivores, potentially favoring herbivores in species rich mixtures (Zhang et al. 2017; Wang et al. 2025) even in regions that are usually considered to feature more specialized herbivore communities (Forister et al. 2015). In contrast, plant pathogens may respond differently to increasing tree species diversity, as the same changes in community composition and leaf traits that reduce host suitability for herbivores may increase transmission opportunities for pathogens or favor susceptible hosts (Schuldt et al. 2017; Rutten et al. 2021). Studies on natural enemies provide an additional perspective, suggesting that higher tree or plant diversity often increases the abundance and diversity of predators and parasitoids and can, in many cases, strengthen top-down control of herbivores, in line with Root (1973) enemies hypothesis (Barnes et al. 2020; Staab & Schuldt 2020; Stemmelen et al. 2022). However, evidence from forests is also mixed and strongly context-dependent in this respect, with tree species richness alone potentially being a poor predictor of changes in predation pressure compared to the structural and functional diversity of the tree community (Staab & Schuldt 2020; Stemmelen et al. 2022). To our knowledge, no biodiversity experiment has yet simultaneously analyzed herbivory, pathogen damage, and predation in relation to tree species richness and extinction scenarios, functional trait structure, and trait diversity, limiting our understanding of how these interactions jointly respond to biodiversity change.

The important role of functional traits and their diversity in mediating effects of biodiversity loss also means that inconsistent effects may be due to the widespread use of random species-loss scenarios in biodiversity experiments. Real-world species extinctions are usually not random with respect to species traits or their positions on an ecological or phylogenetic axis (Purvis et al. 2000; Vamosi & Wilson 2008). For example, rare species and species with small geographic ranges or narrow habitat breadth, as well as species with certain life-history traits, face a higher risk of decline than widespread generalists (Purvis et al. 2000; Suding et al. 2008; Chichorro et al. 2019). Along the leaf economics spectrum, deciduous trees typically show resource-acquisitive leaf traits, e.g., high specific leaf area, low leaf dry matter content and high leaf nutrient concentrations, whereas evergreen trees tend to have tougher leaves, low specific leaf area, high leaf dry matter content and higher C:N ratios (Poorter et al. 2004; Wright et al. 2004; Pringle et al. 2011). Fast-growing and resource-acquisitive strategies are often more vulnerable to drought and other environmental stressors, and accordingly species with such strategies show higher mortality in many systems (Greenwood et al. 2017; Liu et al. 2022). Species extinctions can thus determine the remaining functional diversity of a community in non-random ways. Depending on the driver, extinctions may strongly reduce the occupied trait space and functional differences among the remaining species by removing species with distinct ecological strategies first. In contrast, the loss of functionally similar species (e.g. under intensified competition for limited resources) may leave the occupied trait space largely unchanged and may even increase functional differences among the remaining species, while reducing functional redundancy (Mouillot et al. 2013; Leitão et al. 2016; Weeks et al. 2025). To make species loss experimentally tractable, biodiversity experiments have commonly simulated extinction by removing species according to design rules or along single-trait gradients (Bruelheide et al. 2014). These procedures do not necessarily follow the trait axes that determine real extinction risk (Chen et al. 2020) and may therefore underestimate the effects of non-random, trait-based species loss on multitrophic interactions such as herbivory. Combined trait-based extinction pathways have therefore been proposed as a way to study how non-random species loss moves through ecosystems (Bunker et al. 2005; Chen et al. 2020). However, the consequences of these different trait losses for herbivores, plant pathogens and their natural enemies are still largely unknown.

Here, we use data collected from a large-scale subtropical tree diversity experiment (Bruelheide et al. 2014; Klein et al. 2026) and explicitly combine two complementary approaches. In a first step, we investigate how herbivory and pathogen infestation are influenced by tree species richness and its interplay with leaf trait means and functional diversity across all plots of the experiment. We additionally include predation pressure to test for top-down effects on herbivores. In a second step, we construct explicit extinction pathways by selecting experimental plots that remove tree species along opposing gradients of functional diversity loss (i.e. minimizing and maximizing loss of functional diversity with species richness loss), and compare how these directed species-loss scenarios change the effect of tree species richness on herbivory compared to random species loss. We expect (i) leaf herbivory to increase as tree species richness increases, consistent with greater resource heterogeneity for herbivores, while pathogen infestation shows a negative relationship to tree species richness, due to reduced host concentration and increased barriers to pathogen spread, and (ii) herbivory to be negatively related to predation, consistent with top-down control. We further expect (iii) functional traits of the tree community to drive changes in interaction rates, with herbivory being higher in communities characterized by trait combinations associated with an acquisitive ecological strategy than in more evergreen-dominated communities characterized by resource-conservative strategies, whereas pathogen infestation is expected to be lower in evergreen-dominated and species rich communities. Accordingly, we expect that (iv) effects of tree species richness on herbivory and pathogen infestation depend on the functional composition of the tree community and on non-random species loss. Trait-based species loss will therefore modify trophic interactions more strongly than random loss. We further predict that extinction pathways with strong reductions in functional diversity will weaken tree species richness-driven changes in herbivory and pathogen infestation, whereas extinction pathways which largely retain functional diversity will strengthen richness effects on herbivory and pathogen infestation.

## MATERIAL AND METHODS

### Study site

Our study was conducted at a large-scale forest biodiversity-ecosystem functioning experiment (BEF-China), located in Xingangshan, Dexing, Jiangxi Province, China (29°08′-29°11′N, 117°90′-117°93′E). The climate in this region is classified as subtropical and is characterized by a mean annual temperature of 16.7 °C and a mean annual precipitation of 1821 mm (Yang et al., 2013). The BEF-China experiment comprises a total of 566 experimental plots distributed across two sites, Site A (271 plots, planted in 2009) and Site B (295 plots, planted in 2010). Each plot measures 25.8 × 25.8 m². Within each plot, 400 trees were established in a regular 20 × 20 grid, with species identities randomly drawn from three species pools comprising 42 locally common tree species (Bruelheide et al., 2014). To assess the effects of tree species richness and simulate extinction scenarios, the experiment includes three random extinction scenarios, each based on a ’broken-stick’ model that generates a range of species richness levels (1, 2, 4, 8, 16, and 24 species per plot), as well as two non-random extinction scenarios in which species richness declines systematically by following either local species rarity (rare species lost first) or specific leaf area (SLA, high SLA-species lost first) gradients (Bruelheide et al. 2014).

### Herbivory and pathogen infestation assessment

To quantify levels of herbivory and pathogen damage on planted trees along the species richness gradient, we conducted sampling on 277 plots during September and October 2023, including 140 plots on Site A and 137 plots on Site B. Our methods followed those established in previous studies based on visual estimations of damage (Schuldt et al. 2012; Schuldt et al. 2015). For each selected tree, leaves were sampled from three branches evenly distributed across the canopy, whenever possible. From each branch, 10 leaves were collected, resulting in a total of 30 leaves per tree. Only fully developed leaves from the current annual vegetation period were included in the sampling. The number of trees sampled in each plot varied according to species richness in order to achieve a comprehensive survey of all species present (Table S 1). This ensured a representative sampling of trees across different diversity treatments.

The sampled leaves were visually inspected to estimate the percentage of leaf area missing due to herbivory and the percentage affected by pathogen infestation. These estimates were categorized into predefined damage classes, 0%, 1%-5%, 6%-25%, 26%-50%, 51%-75%, >76% (see also Schuldt et al. 2017). Damage estimates were cross-checked by a second assessor to ensure consistency and reduce observer bias. For each leaf, damage was categorized separately for herbivory and pathogen-related damage and were quantified as mean percentages per tree and aggregated to the tree and plot level. Predefined damage classes were converted to continuous values using mean values per damage class (Schuldt et al. 2015).

### Predation assessment

To measure predation rates, we used artificial caterpillars made from green plasticine (STAEDTLER® Noris® Plasticine green) as proxies for real caterpillars. This method is widely employed to assess predation intensity in ecological studies (Low et al. 2014; Roslin et al. 2017). To quantify predation pressure, we placed artificial caterpillars on two selected trees per plot. Trees were selected as the species with the highest and the lowest previously recorded leaf herbivory among the species present in the plot, based on Schuldt et al. (2017) and unpublished data of 2018 and 2019, to capture the potential within-plot range of predation. On each focal tree, five artificial caterpillars (≈ 2 cm long, ≈ 0.6 cm in diameter; produced with a hand-operated extruder) were attached to the trunk with small needles, resulting in 10 caterpillars per plot. Caterpillars were fixed at 1.4 - 2.0 m height, spaced approximately 10 cm apart, oriented upwards and, where present, placed close to a branch with foliage. The artificial caterpillars were left in place for 14 days, after which they were collected and examined in the laboratory under a microscope. Bite marks and other physical signs of attempted predation or parasitism, such as stings, were recorded. Each caterpillar was assigned a predation status based on the presence or absence of marks (Figure S 1). Predation was therefore calculated as the proportion of artificial caterpillars with visible attack marks per plot.

### Tree species traits, biomass and evenness

Morphological and nutritional leaf functional trait data were obtained from recent measurements in the BEF-China experiment (Davrinche & Haider 2021; Davrinche et al. 2023; Castro Sánchez-Bermejo et al. 2025) and supplemented with entries from the TRY database (Kattge et al. 2020). We first averaged across richness levels to obtain species-level trait means. We then focused on nine leaf traits that capture morphological and nutritional axes of plant strategies and determine leaf quality for primary consumers (Elser et al. 2000; Wright et al. 2004): specific leaf area (SLA), leaf dry matter content (LDMC), leaf carbon (C), nitrogen (N), phosphorus (P), potassium (K), magnesium (Mg), calcium (Ca) concentrations, and the leaf C:N ratio (Table S 2). In addition, we included evergreenness (evergreen vs. deciduous) as a categorical variable to account for differences in community structure and resource-use strategies. Plot-level aboveground biomass for 2023 was derived from field measurements of basal diameter and tree height of individual trees species in each plot and calculated using published allometric equations for BEF-China tree species (Huang et al. 2018).

Shannon diversity and Pielou’s evenness were quantified from species biomass, calculated on a species-by-plot biomass matrix using the R package vegan (Oksanen et al. 2022). The resulting indices were tested as covariates in statistical models.

### Statistical Analysis

All statistical analyses were conducted in R, version 4.3.2 (R-Core-Team 2021). Mean values for herbivory and pathogen infestation were log-transformed (*log1p()*) to improve model residuals.

Whenever trait values were not measured in the field or available from TRY, we imputed missing values with R package missForest (Stekhoven & Stekhoven 2013) using 10,000 trees on a species by trait matrix. Evergreenness, coded as evergreen or deciduous, served as a categorical predictor for the imputation. The out-of-bag error was 0.030 as normalized root mean squared error (NRMSE) for numeric traits and 0.103 as proportion falsely classified (PFC) for the factor variable. In total, 7.7% of trait values were imputed (Table S 3). Evergreenness was used only for imputation and was not part of the functional traits that were used for the functional diversity calculations.

Using species’ relative biomass shares per plot as weights, we computed functional diversity indices with the function dbFD in the FD package (Laliberté et al. 2014). All numeric traits were z-standardized before distance calculations. We calculated Rao’s Q, functional richness, functional evenness, functional dispersion, functional divergence, and community-weighted means (CWMs) of the nine leaf traits stated above. These metrics were used as candidate predictors in subsequent model building (see below). We inspected pairwise correlations among traits and indices and reduced the set of strongly correlated (corr > 6.5) variables to avoid redundancy in subsequent analyses, retaining variables that best represented complementary axes of leaf morphology and nutrition (Figure S 2).

To analyze the effects of predictors on herbivory and pathogen infestation, we fitted a linear mixed-effects model with a Gaussian error structure for each of the two response variables using glmmTMB (Brooks et al. 2017). We started from an initial global model including all uncorrelated variables (predation rate, extinction scenario, tree species richness (log₂-transformed), evenness, evergreenness of the individual tree, plot-level tree species biomass (log₂-transformed), topographic covariates (slope and aspect of the plot), and species specific traits (LDMC, C, N, C:N) or alternative CWM values of those traits and functional-diversity metrics. Random intercepts for tree species and for plots nested within sites (1∣species)+(1∣SITE/PLOT) were included to account for the hierarchical sampling design. We also evaluated a set of ecologically motivated interaction terms: Tree species richness * Evergreenness; Extinction scenarios * Tree species richness; Rao’s Q * Tree species richness; mean predation * Extinction scenarios; mean predation * Tree species richness; mean predation * Rao’s Q; mean predation * Tree species richness * Extinction scenarios; Extinction scenarios * Tree species richness * Evergreenness. Fixed effects were simplified from a global model using AIC-based comparisons of nested models via backward elimination. Models containing Rao’s Q performed as well as or better than models with functional dispersion and had the advantage that the full dataset including monocultures could be used. Thus, Rao’s Q was retained as the chosen FD metric.

In addition to including predation as a predictor of herbivory, we modelled plot-level predation as a response variable in a separate analysis. Predation was quantified as the number of attacked caterpillars out of 10 individuals exposed per plot and analysed using a beta-binomial GLMM fitted with glmmTMB. The model included site, extinction scenario, topography (aspect and slope), tree species richness, and functional diversity (Rao’s Q) as fixed effects, and we additionally tested the Rao’s Q * tree richness interaction. All model assumptions were evaluated using residual diagnostics based on simulated residuals using the R package DHARMa (Hartig 2016), and we checked for overdispersion and singular fits. Post hoc comparisons were conducted using the R package emmeans (Lenth 2023). Model predictions and observed data were visualized using ggplot2 (Wickham 2016).

### Trait-based extinction pathways based on functional diversity

In a second step, we explored how two additional extinction pathways based on functional similarity modify the relationship between tree species richness and herbivory. These pathways were designed to complement the non-random BEF-China scenarios based on rarity and SLA, which represent directed loss along single trait axes. Following the approach by Chen et al. (2020), species were sequentially removed from the regional species pool based on functional diversity (Figure 1). We first calculated a functional distance matrix among all tree species using Gower distances, package FD (Laliberté et al. 2014), on the standardized, multivariate leaf trait matrix (SLA, LDMC, C, N, P, K, Mg, Ca, C:N) . For each species, we then quantified its mean functional distance to all other species. This allowed us to rank species from functionally most distinct to functionally most similar. Based on this ranking, we defined two directed extinction pathways. In the first pathway (FD max), species with the functionally most similar traits were removed first, so that the remaining species pool at a given richness level was dominated by functionally distinct species. In the second pathway (FD min), species with the most distinct traits were removed first, resulting in communities composed of functionally more similar species. For each pathway, five extinction levels were defined based on a species richness of 16, 8, 4, 2, and 1 tree species, by reducing the species pool to 20, 16, 8, 4, and 2 species. Only plots whose species compositions were fully compatible with the reduced pool were retained, ensuring that all scenarios reflected actually observed combinations within the experimental design (Figure S 3). In addition, we generated a set of trait-independent pathways by randomly permuting the species ranking 100 times and applying the same selection procedure, thereby obtaining an ensemble of 100 random extinction pathways. To quantify how these extinction pathways modify the relationship between tree species richness and herbivory as well as pathogen infestation, we fitted linear mixed-effects models for each pathway, with log-transformed herbivory or pathogen damage as the response, log₂-transformed tree species richness as the main fixed effect and random intercepts for tree species and plots.

**Figure 1.**
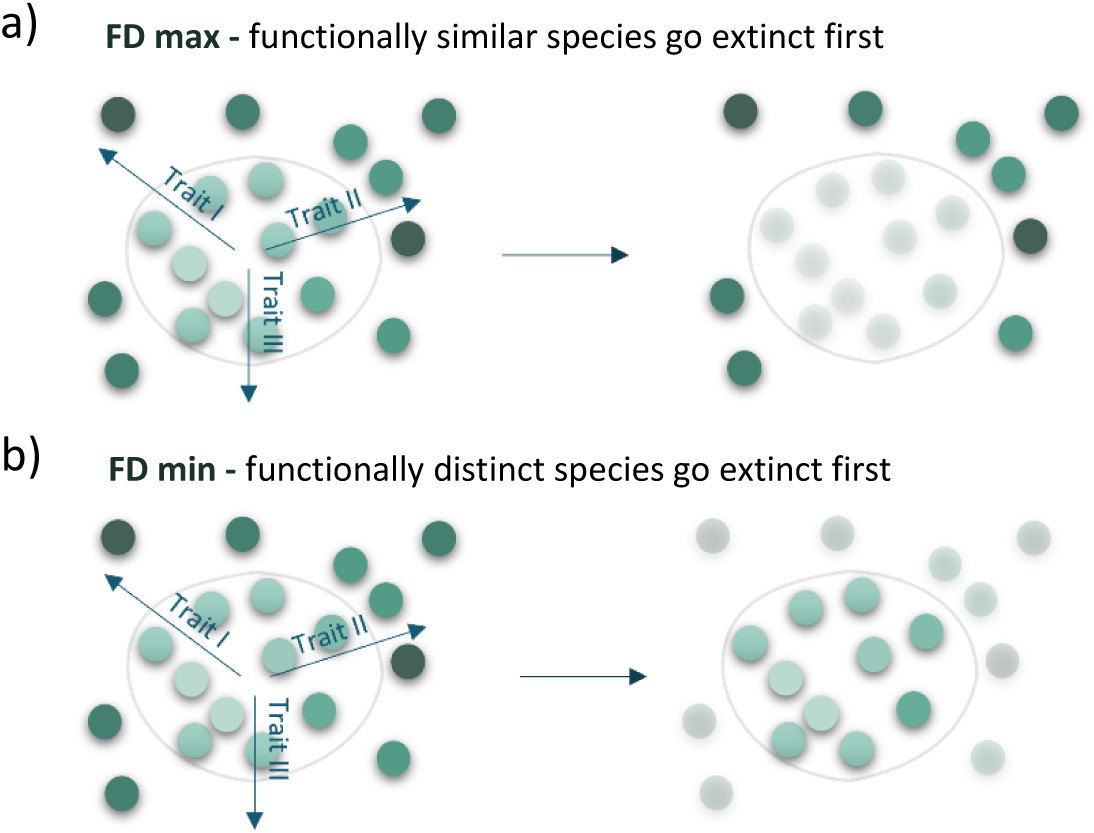
Simplified illustration of the two directed extinction pathways based on functional similarity. Colored circles represent tree species in regional trait space, and the grey outline highlights the species pool retained at a given extinction level. (a) Similar-first pathway: functionally similar species go extinct first, so the remaining pool is dominated by functionally more distinct strategies. (b) Distinct-first pathway: functionally distinct species go extinct first, resulting in a remaining pool composed of more similar strategies.

## RESULTS

### Effects of tree diversity, predation and functional composition on herbivory

Across all tree individuals, mean leaf herbivory was 10.3% (± 8.5 SD), ranging from 0 to 84.2% of leaf area damaged per tree individual. Herbivory was highest on *Diospyros japonica* Siebold & Zucc. and *Idesia polycarpa* Maxim., with average values of 25.0% (± 14.3 SD) and 23.5% (± 16.3 SD) leaf area damaged, and lowest on *Manglietia fordiana* Oliv. with an average of 1.7% (± 2.29 SD).

Models including functional trait CWMs clearly outperformed models based on species-specific trait values (ΔAIC > 10). The final model for log-transformed herbivory included plot-level predation rate, tree species richness, Rao’s Q and CWMs of LDMC and C:N, tree biomass (plot-level per tree species), and evergreenness (individual tree-level), as well as the interactions between predation and extinction scenario and between tree species richness and Rao’s Q (Table 1).

**Table 1.** Final linear mixed-effects model (glmmTMB) testing the relationship of predictors on log-transformed leaf herbivory. Shown are Type II Wald χ² statistics, degrees of freedom (df), P values, estimates for fixed effects, and the standard error (SE). R² marginal = 0.185; R² conditional = 0.494.

| Fixed effect | Estimate | SE | $\chi^2$ | df | P |
| --- | --- | --- | --- | --- | --- |
| Mean predation on plot level | 0.192 | 0.123 | 12.25 | 1 | <0.001 |
| Extinction scenario | - | - | 12.44 | 4 | 0.014 |
| Rao's Q | -0.051 | 0.018 | 2.63 | 1 | 0.105 |
| Tree species richness [$\log_2$] | 0.038 | 0.022 | 9.7 | 1 | 0.002 |
| CWM LDMC | -2.324 | 0.553 | 17.69 | 1 | <0.001 |
| CWM leaf C:N | 0.015 | 0.003 | 16.34 | 1 | <0.001 |
| Tree biomass [log] | 0.008 | 0.006 | 1.73 | 1 | 0.189 |
| Evergreenness | - | - | 25.48 | 1 | <0.001 |
| Predation × Extinction scenario | - | - | 13.26 | 4 | 0.010 |
| Tree species richness [$\log_2$] × Rao's Q | 0.014 | 0.006 | 5.39 | 1 | 0.020 |

Overall, herbivory increased with tree species richness on the log₂ scale, yet the effect depended on the functional diversity of the tree community, as indicated by a significant interaction between log₂ tree species richness and Rao’s Q (Figure 2a). Predation pressure was positively related to herbivory, but this association differed among extinction scenarios as indicated by a significant predation × extinction interaction (Figure 2b, Table 1). Herbivory also differed strongly between evergreen and deciduous communities, with lower predicted herbivory for evergreen individuals than for deciduous trees (Figure 2c). Moreover, herbivory decreased with higher CWMs of LDMC (Figure 2d) and increased with higher CWMs of C:N (Figure 2e). Rao’s Q (as main effect) as well as biomass had no significant effects on log-transformed herbivory (Table 1).

**Figure 2.**
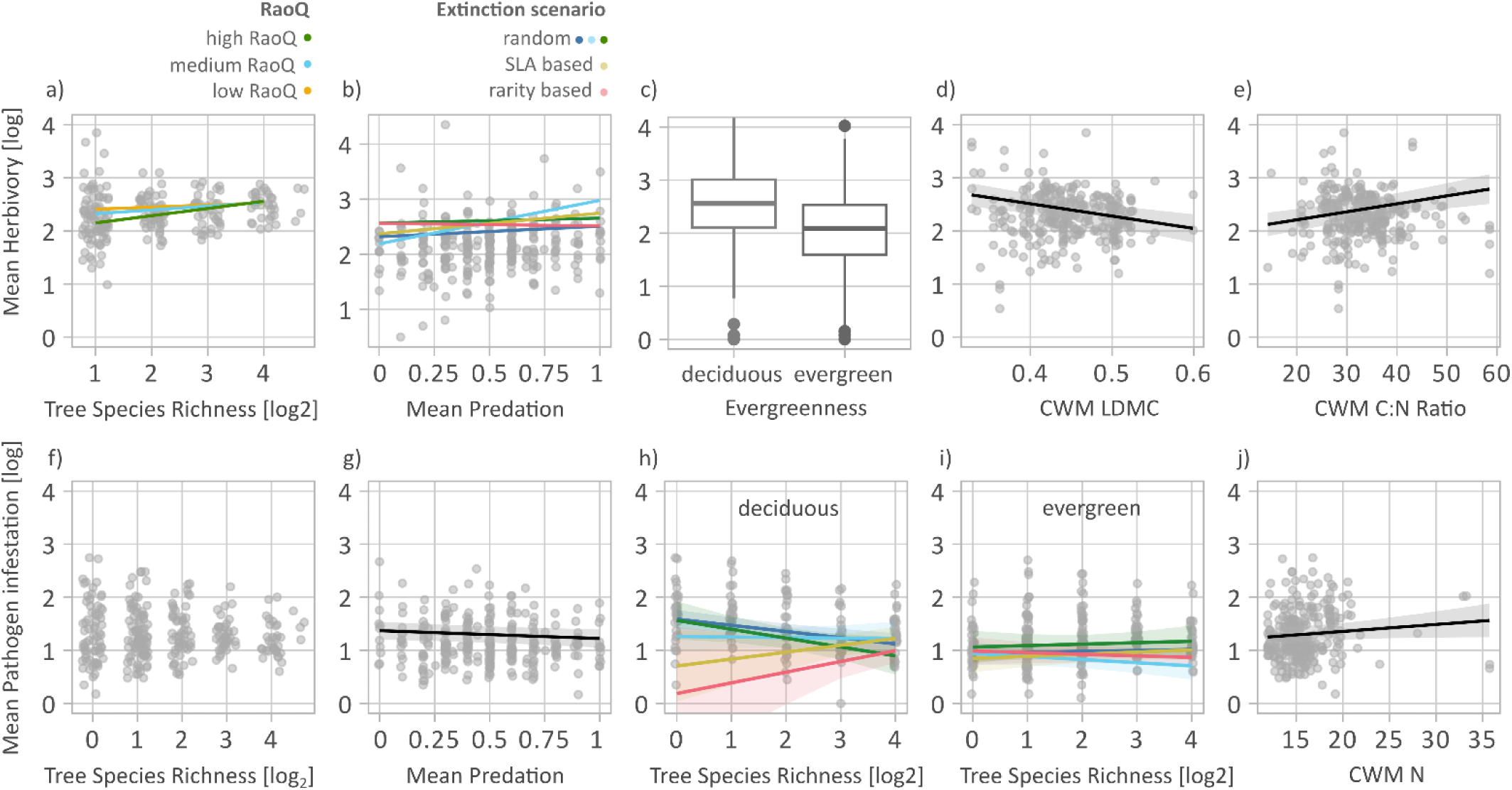
Effect of tree species richness, functional diversity and planted extinction scenarios on mean leaf herbivory (top) and pathogen infestation (bottom). (a) Herbivory (log) and plot-level tree species richness (log₂ scale). Points show plot-level mean herbivory in tree species mixtures, whereas lines show fixed-effect predictions from the mixed model for low, medium, and high values of functional diversity (Rao’s Q). (b) Herbivory (log) and plot-level mean predation. Points show plot-level mean herbivory, whereas lines show fixed-effect predictions from the mixed model for each extinction scenario (three random scenarios vs. directed, trait-based scenarios structured by SLA and rarity). (c) Herbivory (log) in deciduous versus evergreen individuals. (d–e) Herbivory (log) and community-weighted mean (CWM) LDMC (d) and leaf C:N ratio (e). (f) Pathogen infestation (log) and plot-level tree species richness (log₂ scale). (g) Pathogen infestation (log) and plot-level mean predation. (h–i) Pathogen infestation (log) and the number of deciduous (h) and evergreen (i) tree species per plot across extinction scenarios. Lines are colored by extinction-scenario type (three random scenarios vs. trait-based scenarios structured by SLA and rarity). (j) Pathogen infestation (log) and CWM leaf nitrogen content. Outliers with N > 23 were retained in the figure because their exclusion did not change the significance or overall interpretation of the relationship.

### Effects of tree diversity, predation and functional composition on pathogen infestation

Mean pathogen infestation was 2.9% (± 3.9 SD), ranging from 0 to 57.3% per tree. Pathogen infestation was highest on *Ailanthus altissima* and *Nyssa sinensis*, with averages of 8.5% (± 8.45 SD) and 5.7% (± 6.6 SD), and lowest on *Phoebe bournei* with an average of 0.9% (± 1.7 SD). The final model for log-transformed pathogen infestation included mean plot-level predation, extinction scenarios, tree species richness (log₂-transformed), community-weighted mean leaf N, and evergreenness (tree level), as well as their interaction structure extinction scenarios × tree species richness × evergreenness (Table 2).

**Table 2.** Final linear mixed-effects model (glmmTMB) testing the relationship of predictors on log-transformed pathogen infestation. Shown are Type II Wald χ² statistics, degrees of freedom (df), P values, estimates for fixed effects, and the standard error (SE). Marginal R² was 0.075 and conditional R² was 0.353.

| Fixed effect | Estimate | SE | $\chi^2$ | df | P |
| --- | --- | --- | --- | --- | --- |
| Mean predation on plot level | -0.149 | 0.075 | 3.962 | 1 | 0.047 |
| Extinction scenario | - | - | 10.558 | 4 | 0.032 |
| Tree species richness [$\log_2$] | -0.117 | 0.030 | 2.284 | 1 | 0.131 |
| CWM leaf N | 0.013 | 0.008 | 3.006 | 1 | 0.083 |
| Evergreenness | -0.644 | 0.117 | 8.959 | 1 | 0.003 |
| Extinction scenario x Tree species richness [$\log_2$] | - | - | 2.469 | 4 | 0.650 |
| Extinction scenario x Evergreenness | - | - | 10.218 | 4 | 0.037 |
| Tree species richness [$\log_2$] x Evergreenness | 0.134 | 0.033 | 9.873 | 1 | 0.002 |
| Extinction scenario x Tree species richness [$\log_2$] x Evergreenness | - | - | 16.159 | 4 | 0.003 |

Pathogen infestation decreased with increasing predation (Figure 2g). CWM leaf N showed a weak positive association with pathogen infestation (Figure 2i). The Extinction × richness interaction was weak, but the Extinction × evergreenness interaction was supported, indicating scenario-specific shifts between evergreen and deciduous trees (Figure 2h). In addition, the richness × evergreenness interaction was significant, and this dependence of richness effects on evergreenness varied among extinction scenarios, as indicated by the significant three-way interaction.

### Effects of tree diversity and functional composition on predation

Predation on artificial caterpillars averaged 48% (± 25 SD) and ranged from 0% to 100% attacked caterpillars per plot. When modelled as a plot-level response variable, predation showed no evidence for effects of tree species richness, Rao’s Q, or site (Table S3). Instead, predation was weakly associated with topography, with a negative effect of aspect and a positive effect of slope, although both effects were only marginally supported.

### Dissimilarity-based extinction pathways based on functional diversity

In a second step, we used trait-based extinction pathways to investigate how alternative species loss pathways affect the relationship between tree species richness and herbivory or pathogen damage. Across both trait-based extinction pathways, herbivory increased with tree species richness on the log₂ scale, but the strength of this relationship depended on how functional diversity was reduced (Figure 3). Removing functionally similar species first strengthened the richness effect on herbivory compared to removing functionally distinct species, although the relationship remained positive in both cases (functionally similar species removed first: χ² = 23.04, df = 1, P < 0.001; functionally distinct species removed first: χ² = 7.79, df = 1, P = 0.005). Random, trait-independent extinction pathways produced a range of relationships between tree species richness and herbivory. The pathway removing functionally distinct species had a fitted slope of 0.062, which was close to the mean random slope of 0.060 (SD = 0.024, 95% range = 0.005 to 0.101, n = 100), whereas the pathway removing functionally similar species produced a steeper slope of 0.109, exceeding the upper end of the random range. For pathogen infestation, tree species richness decreased infestation in all scenarios (Figure 3). The slope was slightly more negative under the pathway removing functionally distinct species than under the pathway removing functionally similar species first. Specifically, the fitted slope was -0.064 when functionally distinct species were removed first and -0.051 when functionally similar species were removed first. Both values were close to the mean random slope of -0.054 (SD = 0.025, 95% range = -0.105 to -0.009, n = 100).

**Figure 3.**
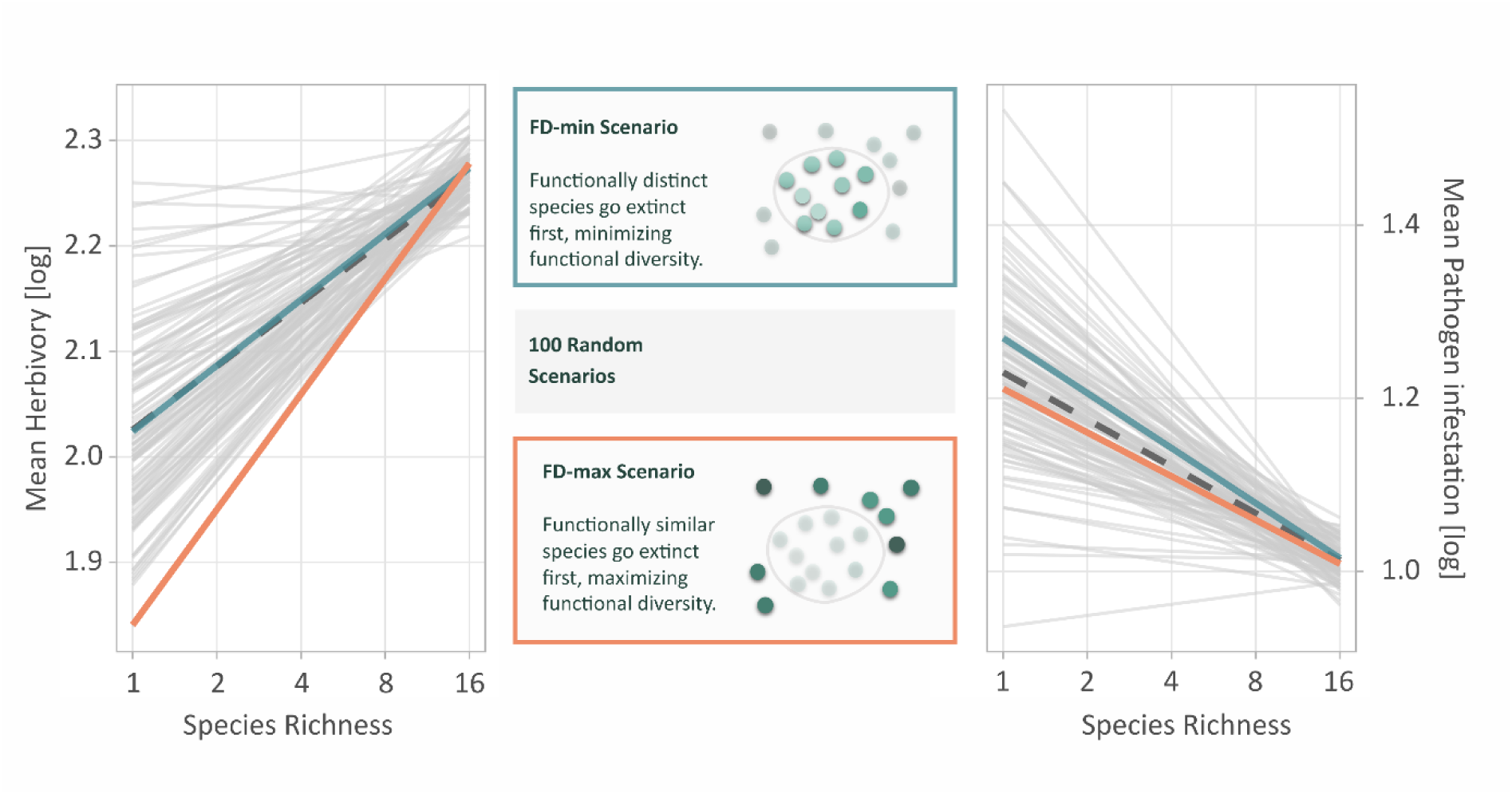
Effects of log₂-transformed tree species richness on log-transformed leaf herbivory (left) and pathogen infestation (right) under different extinction pathways. Grey lines show the range of relationships generated by random, trait-independent extinction pathways (dashed grey line: mean slope across random pathways). The orange line represents the trait-based pathway in which functionally similar tree species were removed first (FD-max; effect of species richness on herbivory: χ² = 20.16, df = 1, P < 0.001). The blue line represents the pathway in which functionally distinct species were removed first (FD-min; effect of species richness on herbivory: χ² = 5.94, df = 1, P = 0.015).

## Discussion

Our findings show that expectations based entirely on the loss of tree species richness may misrepresent both the extent and the early signs of diversity effects on leaf damage, as non-random species loss can alter functional redundancy within the host strategies most relevant to consumers. In herbivory, early loss of functionally similar hosts is associated with a greater decline in herbivory damage as tree species diversity declines, while pathogen responses are likely to remain comparatively weak unless changes in biodiversity also alter host density, spatial continuity, or local infestation conditions. Overall, functional diversity and host redundancy establish the link between biodiversity loss and interaction strength, and thus between changes in tree species richness and shifts in energy and nutrient transfer from trees to higher trophic levels. Our study shows that herbivory increased along with the gradient of tree species richness. Viewed from a biodiversity-loss perspective, this implies that herbivory declines as tree species richness is reduced. This relationship was strongly dependent on the functional diversity of the community, supporting our expectation that tree functional diversity mediates tree richness effects on herbivory. When we introduced non-random, trait-based extinction pathways, herbivory deviated from expectations under random loss. In particular, when functionally similar and thus redundant tree species were removed first, herbivory heavily declined, consistent with our prediction that non-random loss can modify richness effects by changing functional redundancy. In comparison, pathogen infestation showed only weak differences between extinction pathways, providing limited support for a consistently negative tree species richness and pathogen relationship in our system. Predation showed no clear correlation with tree species diversity, and its connection with herbivory was not significant, so there was no evidence of a uniform top-down effect on herbivory, as we had originally expected.

### Functional diversity, tree species richness and extinction pathways

The impact of tree species richness on arthropod herbivory in our study depended on functional diversity. Herbivory decreased with decreasing tree species richness, and this decrease was significantly stronger in communities with higher functional diversity of leaf traits. This interaction between species richness and functional diversity shows that the effects of tree species richness are not simply a matter of having more species, but depend on the functional diversity (here: Rao’s Q) of the community. Accordingly, the negative effect of tree species loss on herbivory was most pronounced in particularly species-poor but functionally diverse communities. A given level of tree species richness can therefore lead to very different outcomes in leaf damage, depending on whether it consists of functionally similar trees (low Rao’s Q of leaf traits) or tree species with a broader mix of leaf traits (high Rao’s Q). Previous synthesis and meta-analytical studies suggest that herbivore responses depend on neighborhood context and herbivore diet breadth, and that increasing leaf trait heterogeneity can modify associational effects depending on herbivore specialization and host configuration (Castagneyrol et al. 2013; Kambach et al. 2016; Jactel et al. 2021). These neighborhood effects are commonly discussed as associational resistance versus associational susceptibility, and their consequences for tree performance can depend on functional traits across forest biodiversity experiments (Jactel & Brockerhoff 2007; Li et al. 2025). Studies from BEF China indicate a dominance of generalists in the community composition of herbivorous arthropods (Zhang et al. 2017; Wang et al. 2026), suggesting an important role of arthropod specialization and available host plants. Accordingly, two non-mutually exclusive mechanisms may explain why higher functional diversity strengthens the relationship between tree species richness and herbivory in our system. First, in communities with generally low Rao’s Q, tree species are functionally more similar and thus more substitutable from the perspective of generalist herbivores. As tree species richness declines, a relatively large share of the remaining species can still fall within the usable resource spectrum of generalists, buffering the reduction in effective host availability and limiting declines in herbivory. By contrast, in communities with high Rao’s Q, herbivory may be boosted at high richness by diet mixing and complementary feeding, but richness loss can more rapidly remove these complementary resources, leading to a steeper decline in herbivore performance and damage (Bernays et al. 1994; Unsicker et al. 2008). Second, higher Rao’s Q may reduce functional continuity among remaining hosts as richness declines, limiting efficient host switching and spillover by generalist herbivores and thereby causing a stronger drop in cumulative herbivory damage (Barbosa et al. 2009; Hahn & Orrock 2016).

Trait based extinction pathway analyses support this by varying the functional diversity of species loss. Herbivory deviated most strongly from expectations under random loss when functionally similar species were removed first, whereas removing functionally distinct species resulted in tree species richness effects on herbivory that largely remained within the random loss range. Looking more broadly at forests dominated by generalist herbivores, our results are consistent with evidence that host functional composition can structure plant herbivore networks (Wang et al. 2020; Wang et al. 2026). By removing functionally distinct species first, the extremes of the trait space are primarily truncated, while leaving a more overlapping set of common host strategies, so that the suitable resource subset changes less and tree species richness effects remain closer to random loss expectations. When species loss removes functionally similar hosts early, herbivores lose many of their “alternative” food options at once. This reduces opportunities for diet mixing and can intensify competition on the remaining suitable hosts, so herbivory may decline to a greater extend with decreasing tree species richness than under random loss (Wang et al. 2022).

While some forest herbivore communities contain a high proportion of generalists that can feed on multiple host species (Bernays et al. 1994; Novotny et al. 2002; Wang et al. 2026), herbivory is often reduced in forests dominated by specialists, where increasing tree species diversity can reduce damage through lower host concentration and associational resistance (Jactel & Brockerhoff 2007). An interesting implication is thus that the effects of the same extinction pathways may differ across forest systems depending on whether herbivore communities are dominated by generalists or specialists. Together with the significant interaction between tree species richness and Rao’s Q, these pathway-dependent responses support the interpretation that redundancy among similar host strategies mediates how tree species richness translates into herbivory, depending on the average host specialization.

In contrast to herbivory, pathogen infestation did not show a uniform response to tree species richness, but varied with the planted extinction scenario and Evergreenness. This contrast is consistent with the idea that pathogen damage is often constrained less by trait heterogeneity of plant communities and more by host association and transmission processes (Hantsch et al. 2014; Schuldt et al. 2017; Rutten et al. 2021). Since pathogens tend to be highly host specific, infection success depends strongly on host identity and density-dependent spread. Although broader-host-range pathogens may occur or have stronger effects in deciduous communities if these species are, on average, less well defended. Rare tree species may also be particularly vulnerable to such pathogens, so that an increase in rare, deciduous tree species could contribute to higher infection.

Increasing tree species richness can therefore reduce effective host density and spatial continuity, resulting in a modest negative tree species richness signal, even when pathogen diversity increases (Hantsch et al. 2014; Keesing & Ostfeld 2021; Rutten et al. 2021). Under such dynamics, changing trait dissimilarities among hosts, as in the constructed extinction pathways, may not translate into large shifts in damage unless it also changes host density, spatial aggregation, or microenvironmental conditions relevant for infection and sporulation, including neighbor identity effects and stand microclimate (Rutten et al. 2021; Saadani et al. 2021; Field et al. 2025).

In this context, the comparatively weak extinction pathway effects on pathogen infestation provide a useful contrast, consistent with the fact that pathogens are often more restricted by host identity than herbivorous arthropods, suggesting that both specialized herbivores and specialized pathogens provide an interesting comparison.

### Top-down control

Predation showed no clear relations with tree species richness or functional diversity. With herbivory as a response, mean predation was positively associated with herbivory, but the strength of this relation differed slightly between the various extinction scenarios. Taken together, these results provide no evidence of a consistent top-down control in herbivory at a plot level. Instead, they are consistent with a primary bottom-up link, in which predator activity follows prey availability and energy flow through the herbivore trophic pathway (Buzhdygan et al. 2020). This interpretation is also supported by evidence that biodiversity effects can cascade bottom-up but are dampened across trophic levels (Scherber et al. 2010). Accordingly, a previous study in BEF-China showed that higher tree species diversity can increase the diversity and abundance of both herbivores and natural enemies, and that predators and parasitoids can enhance tree productivity through top-down control of herbivores, although these effects may vary seasonally (Li et al. 2023). In other words, predator activity can respond to plant diversity through changes in prey communities as well as through changes in habitat structure and microenvironment, which is consistent with prey tracking and a shared driver interpretation (Chen et al. 2023). Another possible explanation is that predator effects on herbivory do exist, but are difficult to detect using our plot-level approach. Predation assessments using artificial caterpillars provide a standardized index of predator attack activity, based on the proportion of models bearing attack marks after a fixed exposure period, and they only capture predator groups that interact with the models and leave identifiable traces (Low et al. 2014; Lövei & Ferrante 2017; Ferrante & Lövei 2025). Aggregation at the plot level can also mask the heterogeneity of predation risk at the tree level, and a discrepancy between the timing of predation assessment and the accumulation of herbivory can decouple current attack rates from seasonal total damage. At the same time, artificial caterpillars detected strong ecological signals in other contexts, including clear responses of predation activity to canopy context and vegetation structure, and in some systems along diversity gradients (Leles et al. 2017; Volf et al. 2022). Against this background, the positive association between predation and herbivory in our model, and its variation among extinction scenarios, is more consistent with predators tracking prey availability and energy flux than with a single, predictable top-down suppression pattern at the plot scale.

### Additional trait effects and why they matter

Because tree species richness can remain unchanged while community composition shifts, considering additional trait axes is important for separating effects of species number from effects of the dominant plant traits or ecological strategies that directly mediate consumer and pathogen interactions. Evergreenness was one of the strongest predictors of leaf damage in our models and points to a compositional axis that is not captured by tree species richness or Rao’s Q alone. Herbivory was consistently lower on evergreen than on deciduous trees, which matches expectations from the leaf economics spectrum (Wright et al. 2004). Evergreen species tend to invest more in leaf longevity and structural protection, which is commonly linked to tougher tissues and lower nutrient concentrations per unit mass, and these properties can reduce accessibility of nutrients to chewing herbivores (Coley & Barone 1996; Wright et al. 2004; Díaz et al. 2016). Notably, pathogen infestation additionally showed a tree species richness and evergreenness interaction, suggesting that richness effects on pathogen attack depend on this compositional axis and reinforcing that trait-mediated shifts can modify diversity interaction relationships beyond tree species richness alone (Schuldt et al. 2017; Rutten et al. 2021).

Community weighted mean traits further clarified which aspects of leaf strategies were linked to herbivory. Herbivory decreased with higher community weighted mean leaf dry matter content, which is consistent with higher tissue density being associated with greater leaf toughness and mechanical resistance that can reduce feeding rates and performance of chewing herbivores (Hanley et al. 2007; Clissold et al. 2009; Onoda et al. 2011). This pattern is broadly consistent with the strong evergreenness signal in our models, because evergreen-dominated communities tend to be characterized by tougher, higher-LDMC leaves and more conservative resource-use strategies. By contrast, herbivory increased with higher community weighted mean C:N ratio. Although a higher C:N ratio often indicates lower nitrogen availability for nitrogen limited herbivores, a positive relationship with herbivory is consistent with compensatory feeding, whereby herbivores increase consumption on nitrogen poor foliage to meet nutrient demands (Mattson 1980; Lavoie & Oberhauser 2004; Behmer 2009). Leaf dry matter content and C:N ratio can covary along conservative strategies, for example when denser leaves are associated with lower nitrogen concentrations (Wright et al. 2004; Onoda et al. 2011; Perez-Harguindeguy et al. 2016). At the same time, the two traits capture partially overlapping but mechanistically distinct constraints. Leaf dry matter content is a proxy for tissue density and is often associated with leaf toughness and mechanical resistance (Perez-Harguindeguy et al. 2016), whereas C:N ratio is commonly interpreted as indicating nitrogen availability and stoichiometric constraints from the perspective of herbivores (Mattson 1980). Their relationships with herbivory may therefore differ even within broadly similar resource use strategies. In contrast to these structural and stoichiometric links to herbivory, pathogen infestation was associated with community weighted mean leaf nitrogen, consistent with infection being constrained by host identity and tissue properties that influence nutrient availability and within-leaf growth conditions (Hantsch et al. 2014; Rutten et al. 2021; Saadani et al. 2021). Interestingly, community weighted means were more informative than species-level trait values in our analyses. This may indicate that plot-level functional composition better reflects the resource environment experienced by herbivore assemblages and the integration of damage across multiple hosts, whereas species-level trait data in our dataset were limited in trait coverage and temporal matching to herbivory. Hence, this supports interpreting both herbivory and pathogen infestation primarily through the functional and compositional context of the surrounding tree community, rather than through species specific trait values alone.

Together, these observed trait associations suggest that herbivory and pathogen infestation respond to the same compositional gradients via partly different mechanisms, which may help explain the weaker and partly opposite pathogen responses along the tree species richness gradient.

## Conclusion

Directed species loss can shift interaction strengths, highlighting a broader implication of our study. Biodiversity loss is likely to restructure energy and nutrient transfer through forest food webs, because different consumer guilds can respond differently to the same compositional change. This helps explain why general statements based solely on tree species richness cannot be generalized across interaction types, and highlights functional diversity and host redundancy as key links between biodiversity loss and shifts in trophic interactions. A key implication is therefore that extinction simulations will be most informative when they are linked more closely to real-world mortality processes and when their consequences are evaluated in a multitrophic framework. Future work could parameterize extinction pathways using empirical risk indicators for forests, such as rarity, drought sensitivity, susceptibility to pests and pathogens, observed mortality patterns, or trait syndromes associated with climatic change, and then test whether these pathways consistently alter interaction strengths across sites, years and environmental contexts. Extending these scenarios to additional tree-associated trophic groups, as well as to ecosystem multifunctionality, would help reveal how non-random biodiversity loss cascades through forest food webs. Combining such extinction pathway designs with approaches that quantify interaction strengths and energy fluxes across trophic levels would strengthen mechanistic forecasts of how ongoing biodiversity decline reshapes forest ecosystem functioning.

From a management perspective, our results suggest that in species-poor stands, mixtures with low redundancy among suitable host strategies can support lower herbivory. Whether lower herbivory is desirable depends on the management objective. In productive stands, moderate herbivory may be compatible with high biomass production, whereas management is typically concerned with preventing persistent or severe damage that compromises growth and yield. In this sense, functional redundancy provides a trait-based tool for designing forest tree species mixtures. Overall, our results reinforce the value of maintaining tree species-rich forests, because higher diversity not only supports multiple ecosystem functions but also reduces the likelihood that non-random species loss will cause unexpected shifts in trophic interactions.

## Acknowledgements

We gratefully acknowledge the support of the BEF-China platform and the Zhejiang Qianjiang-yuan Forest Biodiversity National Observation and Research Station (QForDiv). Our sincere appreciation goes to Helge Bruelheide, Keping Ma, and Bernhard Schmid for initiating the BEF-China platform, and to Xiaojuan Liu for her role in managing the platform. We extend our gratitude to Alexandra-M. Klein and Chao-Dong Zhu for their combined efforts in coordinating the research unit MultiTroph (452861007/FOR 5281). We acknowledge Shan Li and Bo Yang for their assistance in maintaining the research station. Further we want to thank all the passionate student helpers that have been involved in the field work, Nina Donhauser, Thomas Hruschka, Dilixiati Kaiwusaier, Lea Osterloh, Florian Rosenthal, Nina Stempel, Johannes Wawrzinek and Laura Zielinsky. Pablo Castro Sánchez-Bermejo, Andréa Davrinche, and Sylvia Haider were supported by the International Research Training Group TreeDì, jointly funded by the Deutsche Forschungsgemeinschaft (DFG, German Research Foundation) 319936945/GRK2324 and the University of Chinese Academy of Sciences (UCAS).

## Author contributions

Mareike Mittag collected the data, compiled the dataset and led the writing of the manuscript. Mareike Mittag analyzed the data with support from Georg Albert and advice from Ming-Qi-ang Wang. Pablo Castro Sánchez-Bermejo, Andréa Davrinche, Sylvia Haider, Shan Li and Xiaojuan Liu contributed data. Georg Albert, Andreas Schuldt and Jana S. Petermann contributed to interpretation of the results and critically revised the manuscript. All authors have reviewed and commented on the manuscript.

## Data availability statement

Data will be made available upon submission via the BEF-China project database (https://data.botanik.uni-halle.de/bef-china/).

## Conflict of interest statement

The authors have no conflict of interest to declare.

## Notes

### Competing Interest Statement

The authors have declared no competing interest.

## References

1. Barbosa, P., Hines, J., Kaplan, I., Martinson, H., Szczepaniec, A. & Szendrei, Z. (2009). Associational resistance and associational susceptibility: having right or wrong neighbors. Annual review of ecology, evolution, and systematics, 40, 1–20.

2. Barnes, A.D., Scherber, C., Brose, U., Borer, E.T., Ebeling, A., Gauzens, B. et al. (2020). Biodiversity enhances the multitrophic control of arthropod herbivory. Science Advances, 6, eabb6603.

3. Behmer, S.T. (2009). Insect herbivore nutrient regulation. Annual review of entomology, 54, 165–187.

4. Bernays, E., Bright, K., Gonzalez, N. & Angel, J. (1994). Dietary mixing in a generalist herbivore: tests of two hypotheses. Ecology, 75, 1997–2006.

5. Brooks, M.E., Kristensen, K., Van Benthem, K.J., Magnusson, A., Berg, C.W., Nielsen, A., et al. (2017). glmmTMB balances speed and flexibility among packages for zero-inflated generalized linear mixed modeling.

6. Bruelheide, H., Nadrowski, K., Assmann, T., Bauhus, J., Both, S., Buscot, F. et al. (2014). Designing forest biodiversity experiments: general considerations illustrated by a new large experiment in subtropical China. Methods in Ecology and Evolution, 5, 74–89.

7. Bunker, D.E., DeClerck, F., Bradford, J.C., Colwell, R.K., Perfecto, I., Phillips, O.L. et al. (2005). Species loss and aboveground carbon storage in a tropical forest. Science, 310, 1029–1031.

8. Buzhdygan, O.Y., Meyer, S.T., Weisser, W.W., Eisenhauer, N., Ebeling, A., Borrett, S.R. et al. (2020). Biodiversity increases multitrophic energy use efficiency, flow and storage in grasslands. Nature Ecology & Evolution, 4, 393–405.

9. Castagneyrol, B., Giffard, B., Péré, C. & Jactel, H. (2013). Plant apparency, an overlooked driver of associational resistance to insect herbivory. Journal of Ecology, 101, 418–429.

10. Castro Sánchez-Bermejo, P., Carmona, C.P., Schuman, M.C., Benavides, R., Sachsenmaier, L., Li, S., et al. (2025). Intraspecific and intraindividual trait variability decrease with tree richness in a subtropical tree biodiversity experiment. Nature Communications, 16, 11009.

11. Chen, J.-T., Wang, M.-Q., Li, Y., Chesters, D., Luo, A., Zhang, W. et al. (2023). Functional and phylogenetic relationships link predators to plant diversity via trophic and non-trophic pathways. Proceedings of the Royal Society B: Biological Sciences, 290.

12. Chen, Y., Huang, Y., Niklaus, P.A., Castro-Izaguirre, N., Clark, A.T., Bruelheide, H. et al. (2020). Directed species loss reduces community productivity in a subtropical forest biodiversity experiment. Nature Ecology & Evolution, 4, 550–559.

13. Chichorro, F., Juslén, A. & Cardoso, P. (2019). A review of the relation between species traits and extinction risk. Biological Conservation, 237, 220–229.

14. Clissold, F.J., Sanson, G.D., Read, J. & Simpson, S.J. (2009). Gross vs. net income: how plant toughness affects performance of an insect herbivore. Ecology, 90, 3393–3405.

15. Coley, P.D. & Barone, J. (1996). Herbivory and plant defenses in tropical forests. Annual review of ecology and systematics, 27, 305–335.

16. Cowie, R.H., Bouchet, P. & Fontaine, B. (2022). The Sixth Mass Extinction: fact, fiction or speculation? Biological Reviews, 97, 640–663.

17. Davrinche, A., Bittner, A., Bruelheide, H., Albert, G., Harpole, W.S. & Haider, S. (2023). High within-tree leaf trait variation and its response to species diversity and soil nutrients. bioRxiv, 2023.2003. 2008.531739.

18. Davrinche, A. & Haider, S. (2021). Intra-specific leaf trait responses to species richness at two different local scales. Basic and Applied Ecology, 55, 20–32.

19. Díaz, S., Kattge, J., Cornelissen, J.H., Wright, I.J., Lavorel, S., Dray, S. et al. (2016). The global spectrum of plant form and function. Nature, 529, 167–171.

20. Dirzo, R., Young, H.S., Galetti, M., Ceballos, G., Isaac, N.J. & Collen, B. (2014). Defaunation in the Anthropocene. science, 345, 401–406.

21. Dunne, J.A., Williams, R.J. & Martinez, N.D. (2002). Network structure and biodiversity loss in food webs: robustness increases with connectance. Ecology letters, 5, 558–567.

22. Elser, J.J., Fagan, W.F., Denno, R.F., Dobberfuhl, D.R., Folarin, A., Huberty, A. et al. (2000). Nutritional constraints in terrestrial and freshwater food webs. Nature, 408, 578–580.

23. Ferrante, M. & Lövei, G.L. (2025). The sentinel approach to quantify ecosystem function intensities. Methods in Ecology and Evolution, 16, 2305–2317.

24. Field, E., Hector, A., Barsoum, N. & Koricheva, J. (2025). Tree diversity reduces pathogen damage in temperate forests: A systematic review and meta-analysis. Forest Ecology and Management, 578, 122398.

25. Forister, M.L., Novotny, V., Panorska, A.K., Baje, L., Basset, Y., Butterill, P.T. et al. (2015). The global distribution of diet breadth in insect herbivores. Proceedings of the National Academy of Sciences, 112, 442–447.

26. Greenwood, S., Ruiz-Benito, P., Martínez-Vilalta, J., Lloret, F., Kitzberger, T., Allen, C.D. et al. (2017). Tree mortality across biomes is promoted by drought intensity, lower wood density and higher specific leaf area. Ecology letters, 20, 539–553.

27. Grossman, J.J., Cavender-Bares, J., Reich, P.B., Montgomery, R.A. & Hobbie, S.E. (2019). Neighborhood diversity simultaneously increased and decreased susceptibility to contrasting herbivores in an early stage forest diversity experiment. Journal of Ecology, 107, 1492–1505.

28. Grossman, J.J., Vanhellemont, M., Barsoum, N., Bauhus, J., Bruelheide, H., Castagneyrol, B. et al. (2018). Synthesis and future research directions linking tree diversity to growth, survival, and damage in a global network of tree diversity experiments. Environmental and Experimental Botany, 152, 68–89.

29. Grünig, M., Mazzi, D., Calanca, P., Karger, D.N. & Pellissier, L. (2020). Crop and forest pest metawebs shift towards increased linkage and suitability overlap under climate change. Communications Biology, 3, 233.

30. Hahn, P.G. & Orrock, J.L. (2016). Neighbor palatability generates associational effects by altering herbivore foraging behavior. Ecology, 97, 2103–2111.

31. Hanley, M.E., Lamont, B.B., Fairbanks, M.M. & Rafferty, C.M. (2007). Plant structural traits and their role in anti-herbivore defence. Perspectives in Plant Ecology, Evolution and Systematics, 8, 157–178.

32. Hantsch, L., Bien, S., Radatz, S., Braun, U., Auge, H. & Bruelheide, H. (2014). Tree diversity and the role of non-host neighbour tree species in reducing fungal pathogen infestation. Journal of Ecology, 102, 1673–1687.

33. Hartig, F. (2016). DHARMa: residual diagnostics for hierarchical (multi-level/mixed) regression models. CRAN: Contributed Packages.

34. Hooper, D.U., Adair, E.C., Cardinale, B.J., Byrnes, J.E., Hungate, B.A., Matulich, K.L. et al. (2012). A global synthesis reveals biodiversity loss as a major driver of ecosystem change. Nature, 486, 105–108.

35. Huang, Y., Chen, Y., Castro-Izaguirre, N., Baruffol, M., Brezzi, M., Lang, A. et al. (2018). Impacts of species richness on productivity in a large-scale subtropical forest experiment. Science, 362, 80–83.

36. Isbell, F., Calcagno, V., Hector, A., Connolly, J., Harpole, W.S., Reich, P.B. et al. (2011). High plant diversity is needed to maintain ecosystem services. Nature, 477, 199–202.

37. Jactel, H. & Brockerhoff, E.G. (2007). Tree diversity reduces herbivory by forest insects. Ecology letters, 10, 835–848.

38. Jactel, H., Moreira, X. & Castagneyrol, B. (2021). Tree diversity and forest resistance to insect pests: patterns, mechanisms, and prospects. Annual Review of Entomology, 66, 277–296.

39. Kambach, S., Kuehn, I., Castagneyrol, B. & Bruelheide, H. (2016). The impact of tree diversity on different aspects of insect herbivory along a global temperature gradient-a meta-analysis. PLoS One, 11, e0165815.

40. Kattge, J., Bönisch, G., Díaz, S., Lavorel, S., Prentice, I.C., Leadley, P. et al. (2020). TRY plant trait database–enhanced coverage and open access. Global change biology, 26, 119–188.

41. Keesing, F. & Ostfeld, R.S. (2021). Dilution effects in disease ecology. Ecology letters, 24, 2490–2505.

42. Klein, A.-M., Bruelheide, H., Chesters, D., Diekötter, T., Erfmeier, A., Feldhaar, H. et al. (2026). MultiTroph: Multi-trophic interactions in a forest biodiversity experiment in China. Research Ideas and Outcomes, 12, e181743.

43. Laliberté, E., Legendre, P., Shipley, B. & Laliberté, M.E. (2014). Package ‘fd’. Measuring functional diversity from multiple traits, and other tools for functional ecology, 1, 0–12.

44. Langellotto, G.A. & Denno, R.F. (2004). Responses of invertebrate natural enemies to complex-structured habitats: a meta-analytical synthesis. Oecologia, 139, 1–10.

45. Lavoie, B. & Oberhauser, K.S. (2004). Compensatory feeding in Danaus plexippus (Lepidoptera: Nymphalidae) in response to variation in host plant quality. Environmental entomology, 33, 1062–1069.

46. Leitão, R.P., Zuanon, J., Villéger, S., Williams, S.E., Baraloto, C., Fortunel, C. et al. (2016). Rare species contribute disproportionately to the functional structure of species assemblages. Proceedings of the Royal Society B: Biological Sciences, 283.

47. Leles, B., Xiao, X., Pasion, B.O., Nakamura, A. & Tomlinson, K.W. (2017). Does plant diversity increase top-down control of herbivorous insects in tropical forest? Oikos, 126, 1142–1149.

48. Lenth, R. (2023). emmeans: Estimated Marginal Means, aka Least-Squares Means_. R package version 1.8. 5.

49. Li, Y., Schmid, B., Schuldt, A., Li, S., Wang, M.-Q., Fornoff, F. et al. (2023). Multitrophic arthropod diversity mediates tree diversity effects on primary productivity. Nature Ecology & Evolution, 1–9.

50. Li, Y., Schuldt, A., Bauhus, J., Belluau, M., Berthelot, S., Burghardt, K.T. et al. (2025). The tree growth–herbivory relationship depends on functional traits across forest biodiversity experiments. Nature Ecology & Evolution, 9, 2014–2024.

51. Liu, X., Huang, Y., Chen, L., Li, S., Bongers, F.J., Castro-Izaguirre, N. et al. (2022). Species richness, functional traits and climate interactively affect tree survival in a large forest biodiversity experiment. Journal of Ecology, 110, 2522–2531.

52. Lövei, G.L. & Ferrante, M. (2017). A review of the sentinel prey method as a way of quantifying invertebrate predation under field conditions. Insect Science, 24, 528–542.

53. Low, P.A., Sam, K., McArthur, C., Posa, M.R.C. & Hochuli, D.F. (2014). Determining predator identity from attack marks left in model caterpillars: guidelines for best practice. Entomologia Experimentalis et Applicata, 152, 120–126.

54. Mattson, W.J. (1980). Herbivory in relation to plant nitrogen content. Annual review of ecology and systematics, 11, 119–161.

55. Michalko, R., Songsangchote, C., Saksongmuang, V., Wongprom, P., Trisurat, Y. & Košulič, O. (2024). Transformation of dry dipterocarp to dry evergreen forests alters food webs of web-building spiders and their prey. Journal of Insect Conservation, 28, 1363–1373.

56. Mouillot, D., Bellwood, D.R., Baraloto, C., Chave, J., Galzin, R., Harmelin-Vivien, M. et al. (2013). Rare species support vulnerable functions in high-diversity ecosystems. PLoS biology, 11, e1001569.

57. Novotny, V., Basset, Y., Miller, S.E., Weiblen, G.D., Bremer, B., Cizek, L. et al. (2002). Low host specificity of herbivorous insects in a tropical forest. Nature, 416, 841–844.

58. Oksanen, J., Simpson, G., Blanchet, F., Kindt, R., Legendre, P., Minchin, P. et al. (2022). Vegan: Community Ecology Package (R Package Version 2.6-2). 2022.

59. Onoda, Y., Westoby, M., Adler, P.B., Choong, A.M., Clissold, F.J., Cornelissen, J.H. et al. (2011). Global patterns of leaf mechanical properties. Ecology letters, 14, 301–312.

60. Perez-Harguindeguy, N., Diaz, S., Garnier, E., Lavorel, S., Poorter, H., Jaureguiberry, P. et al. (2016). Corrigendum to: New handbook for standardised measurement of plant functional traits worldwide. Australian Journal of botany, 64, 715–716.

61. Poorter, L., Van de Plassche, M., Willems, S. & Boot, R. (2004). Leaf traits and herbivory rates of tropical tree species differing in successional status. Plant biology, 6, 746–754.

62. Pringle, E.G., Adams, R.I., Broadbent, E., Busby, P.E., Donatti, C.I., Kurten, E.L. et al. (2011). Distinct leaf-trait syndromes of evergreen and deciduous trees in a seasonally dry tropical forest. Biotropica, 43, 299–308.

63. Purvis, A., Agapow, P.-M., Gittleman, J.L. & Mace, G.M. (2000). Nonrandom extinction and the loss of evolutionary history. Science, 288, 328–330.

64. R-Core-Team (2021). R: A Language and Environment for Statistical Computing. R Foundation for Statistical Computing Vienna, Austria.

65. Root, R.B. (1973). Organization of a plant-arthropod association in simple and diverse habitats: the fauna of collards (Brassica oleracea). Ecological monographs, 43, 95–124.

66. Roslin, T., Hardwick, B., Novotny, V., Petry, W.K., Andrew, N.R., Asmus, A. et al. (2017). Higher predation risk for insect prey at low latitudes and elevations. Science, 356, 742–744.

67. Rutten, G., Hönig, L., Schwaß, R., Braun, U., Saadani, M., Schuldt, A. et al. (2021). More diverse tree communities promote foliar fungal pathogen diversity, but decrease infestation rates per tree species, in a subtropical biodiversity experiment. Journal of Ecology, 109, 2068–2080.

68. Saadani, M., Hönig, L., Bien, S., Koehler, M., Rutten, G., Wubet, T. et al. (2021). Local Tree Diversity Suppresses Foliar Fungal Infestation and Decreases Morphological but Not Molecular Richness in a Young Subtropical Forest. Journal of Fungi, 7, 173.

69. Scherber, C., Eisenhauer, N., Weisser, W.W., Schmid, B., Voigt, W., Fischer, M. et al. (2010). Bottom-up effects of plant diversity on multitrophic interactions in a biodiversity experiment. Nature, 468, 553–556.

70. Schuldt, A., Baruffol, M., Bruelheide, H., Chen, S., Chi, X., Wall, M. et al. (2014). Woody plant phylogenetic diversity mediates bottom–up control of arthropod biomass in species-rich forests. Oecologia, 176, 171–182.

71. Schuldt, A., Bruelheide, H., Durka, W., Eichenberg, D., Fischer, M., Kröber, W. et al. (2012). Plant traits affecting herbivory on tree recruits in highly diverse subtropical forests. Ecology letters, 15, 732–739.

72. Schuldt, A., Bruelheide, H., Härdtle, W., Assmann, T., Li, Y., Ma, K. et al. (2015). Early positive effects of tree species richness on herbivory in a large-scale forest biodiversity experiment influence tree growth. Journal of Ecology, 103, 563–571.

73. Schuldt, A., Ebeling, A., Kunz, M., Staab, M., Guimarães-Steinicke, C., Bachmann, D. et al. (2019). Multiple plant diversity components drive consumer communities across ecosystems. Nature Communications, 10.

74. Schuldt, A., Hönig, L., Li, Y., Fichtner, A., Härdtle, W., von Oheimb, G., et al. (2017). Herbivore and pathogen effects on tree growth are additive, but mediated by tree diversity and plant traits. Ecology and evolution, 7, 7462–7474.

75. Seidl, R., Thom, D., Kautz, M., Martin-Benito, D., Peltoniemi, M., Vacchiano, G. et al. (2017). Forest disturbances under climate change. Nature climate change, 7, 395–402.

76. Staab, M. & Schuldt, A. (2020). The Influence of Tree Diversity on Natural Enemies—a Review of the “Enemies” Hypothesis in Forests. Current Forestry Reports, 6, 243–259.

77. Stekhoven, D.J. & Stekhoven, M.D.J. (2013). Package ‘missForest’. R package version, 1, 21.

78. Stemmelen, A., Jactel, H., Brockerhoff, E. & Castagneyrol, B. (2022). Meta-analysis of tree diversity effects on the abundance, diversity and activity of herbivores’ enemies. Basic and Applied Ecology, 58, 130–138.

79. Suding, K.N., Lavorel, S., Chapin Iii, F., Cornelissen, J.H., Diaz, S., Garnier, E., et al. (2008). Scaling environmental change through the community-level: A trait-based response-and-effect framework for plants. Global Change Biology, 14, 1125–1140.

80. Unsicker, S.B., Oswald, A., Köhler, G. & Weisser, W.W. (2008). Complementarity effects through dietary mixing enhance the performance of a generalist insect herbivore. Oecologia, 156, 313–324.

81. Vamosi, J.C. & Wilson, J.R. (2008). Nonrandom extinction leads to elevated loss of angiosperm evolutionary history. Ecology Letters, 11, 1047–1053.

82. Volf, M., Volfová, T., Seifert, C.L., Ludwig, A., Engelmann, R.A., Jorge, L.R. et al. (2022). A mosaic of induced and non-induced branches promotes variation in leaf traits, predation and insect herbivore assemblages in canopy trees. Ecology letters, 25, 729–739.

83. Wan, N.-F., Wang, Y.-Q., Fu, L., Liu, J., Woodcock, B.A., Hu, Y.-Q. et al. (2026). Global evidence that plant diversity suppresses pests and promotes plant performance and crop production. Nature Ecology & Evolution, 1–15.

84. Wang, M.-Q., Albert, G., Seifert, C.L., Chesters, D., Bruelheide, H., Li, Y. et al. (2026). Asynchrony and functional diversity couple herbivore community dynamics to host plant diversity. Nature Communications.

85. Wang, M.Q., Albert, G., Chesters, D., Bruelheide, H., Li, Y., Chen, J.T. et al. (2025). Tree diversity, tree growth, and microclimate independently structure Lepidoptera herbivore community stability. Ecological Monographs, 95, e70026.

86. Wang, M.Q., Li, Y., Chesters, D., Bruelheide, H., Ma, K., Guo, P.F. et al. (2020). Host functional and phylogenetic composition rather than host diversity structure plant–herbivore networks. Molecular Ecology, 29, 2747–2762.

87. Wang, M.Q., Yan, C., Luo, A., Li, Y., Chesters, D., Qiao, H.J. et al. (2022). Phylogenetic relatedness, functional traits, and spatial scale determine herbivore co-occurrence in a subtropical forest. Ecological Monographs, 92, e01492.

88. Weeks, T.L., Walkden, P.A., Edwards, D.P., Lees, A.C., Pigot, A.L., Purvis, A. et al. (2025). Land-use change undermines the stability of avian functional diversity. Nature.

89. Wein, A., Bauhus, J., Bilodeau-Gauthier, S., Scherer-Lorenzen, M., Nock, C. & Staab, M. (2016). Tree species richness promotes invertebrate herbivory on congeneric native and exotic tree saplings in a young diversity experiment. PLoS One, 11, e0168751.

90. Wickham, H. (2016). Data analysis. In: ggplot2: elegant graphics for data analysis. Springer, pp. 189–201.

91. Wright, I.J., Reich, P.B., Westoby, M., Ackerly, D.D., Baruch, Z., Bongers, F. et al. (2004). The worldwide leaf economics spectrum. nature, 428, 821–827.

92. Zhang, J., Bruelheide, H., Chen, X., Eichenberg, D., Kröber, W., Xu, X. et al. (2017). Tree diversity promotes generalist herbivore community patterns in a young subtropical forest experiment. Oecologia, 183, 455–467.

