## Supplements for "Trait dissimilarity-based tree species loss affects tree diversity effects on herbivory"

**Table S 1.** Sampling effort for herbivory assessments across tree species richness levels.

| Tree species richness level | Number of plots | Minimum number of sampled trees per plot | Number of trees per tree species | Total number of trees |
|---|---|---|---|---|
| **1** | 40 | 4 | 4 | 160 |
| **2** | 44 | 6 | 3 | 264 |
| **4** | 28 | 12 | 3 | 336 |
| **8** | 20 | 16 | 2 | 320 |
| **16** | 16 | 24 | 1-2* | 384 |
| **24** | 2 | 24 | 1 | 48 |
| | | *Total per site:* | | *1512* |
| | | *Total across both sites:* | | *3024* |

Number of plots, minimum number of trees assessed per plot, mean number of trees assessed per tree species, and total number of trees assessed for leaf herbivory at each tree species richness level. Herbivory was quantified on 30 fully developed leaves per tree (10 leaves from each of three canopy branches per tree). The same sampling scheme was applied at both experimental sites, resulting in 1,512 trees assessed per site and 3,024 trees in total.

* To assess the 24 trees, every present tree species got at least sampled once.

**Table S 2.** Species-level leaf traits used for functional diversity and community-weighted mean calculations.

| Trait | Abbreviation | Unit |
|---|---|---|
| Specific leaf area | SLA | $cm^2\ g^{-1}$ |
| Leaf dry matter content | LDMC | $g\ g^{-1}$ |
| Leaf carbon concentration | C | $mg\ g^{-1}$ |
| Leaf nitrogen concentration | N | $mg\ g^{-1}$ |
| Leaf phosphorus concentration | P | $mg\ g^{-1}$ |
| Leaf potassium concentration | K | $mg\ g^{-1}$ |
| Leaf magnesium concentration | Mg | $mg\ g^{-1}$ |
| Leaf calcium concentration | Ca | $mg\ g^{-1}$ |
| Leaf C:N ratio | C:N | – |

**Table S 3.** Overview of missing trait values in the BEF-China dataset, the proportion filled from TRY, and the proportion imputed for each leaf trait.

| Trait | Percent missing in BEF dataset | Percent filled from TRY | Percent imputed |
|---|---|---|---|
| Specific leaf area | 40 | 40 | 0 |
| Leaf dry matter content | 40 | 40 | 0 |
| Leaf carbon concentration | 40 | 40 | 0 |
| Leaf nitrogen concentration | 40 | 40 | 0 |
| Leaf phosphorus concentration | 40 | 27.5 | 12.5 |
| Leaf potassium concentration | 40 | 25 | 15 |
| Leaf magnesium concentration | 40 | 12.5 | 27.5 |
| Leaf calcium concentration | 40 | 12.5 | 27.5 |
| Leaf C:N ratio | 40 | 40 | 0 |

**Table S 4.** Final generalized linear mixed-effects model (glmmTMB) testing the effects of predictors on predation on plot level. Shown are Type II Wald $\chi^2$ statistics, degrees of freedom (df), P values, estimates for fixed effects, and the standard error (SE).

| Fixed effect | $\chi^2$ | df | P | Estimate | SE |
|---|---|---|---|---|---|
| SITE | 0.026 | 1 | 0.873 | - | - |
| Mean aspect | 3.479 | 1 | 0.062 | -0.001 | 0.001 |
| Mean slope | 3.174 | 1 | 0.075 | 0.018 | 0.010 |
| Rao’s Q | 0.043 | 1 | 0.835 | -0.006 | 0.028 |
| Tree species richness [$log_2$] | 1.78 | 1 | 0.182 | 0.079 | 0.059 |

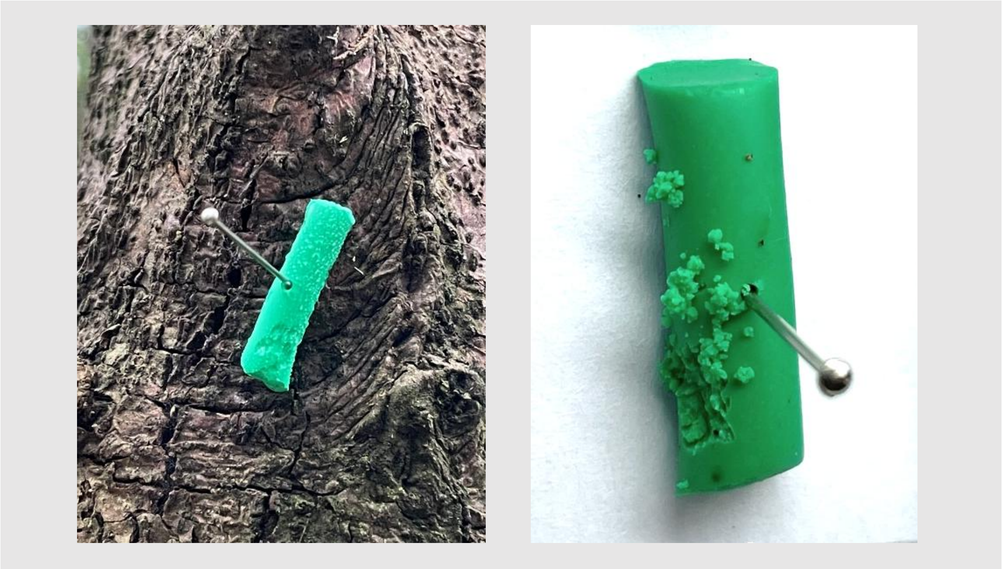

**Figure S 1**. Examples of predation marks on artificial caterpillars.

Bite marks and other physical signs of attempted predation, such as rasping or sting punctures, were recorded on each artificial caterpillar. Caterpillars were classified as “attacked” (predation status = 1) when at least one clear mark was present and as “unattacked” (predation status = 0) otherwise. Predation rates at the plot level were then calculated as the proportion of attacked caterpillars.

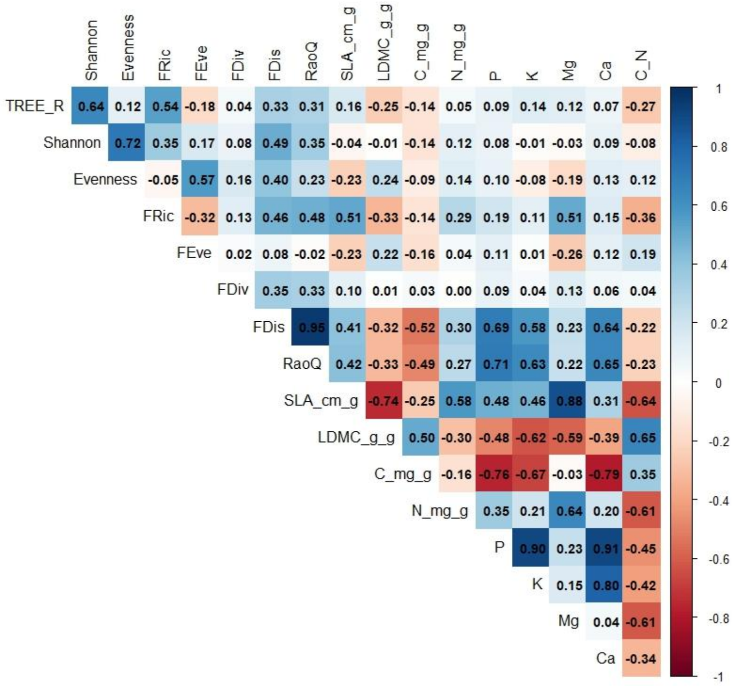


**Figure S 2.** Pairwise correlations among tree community variables used during model building.

Shown are Shannon diversity and Pielou's evenness, functional diversity metrics (FRic, FEve, FDiv, FDis, Rao's Q) and community-weighted mean leaf traits (SLA, LDMC, C, N, P, K, Mg, Ca, and C:N). Colors indicate the direction and strength of correlations (blue = positive, red = negative), and numbers give the correlation coefficients.

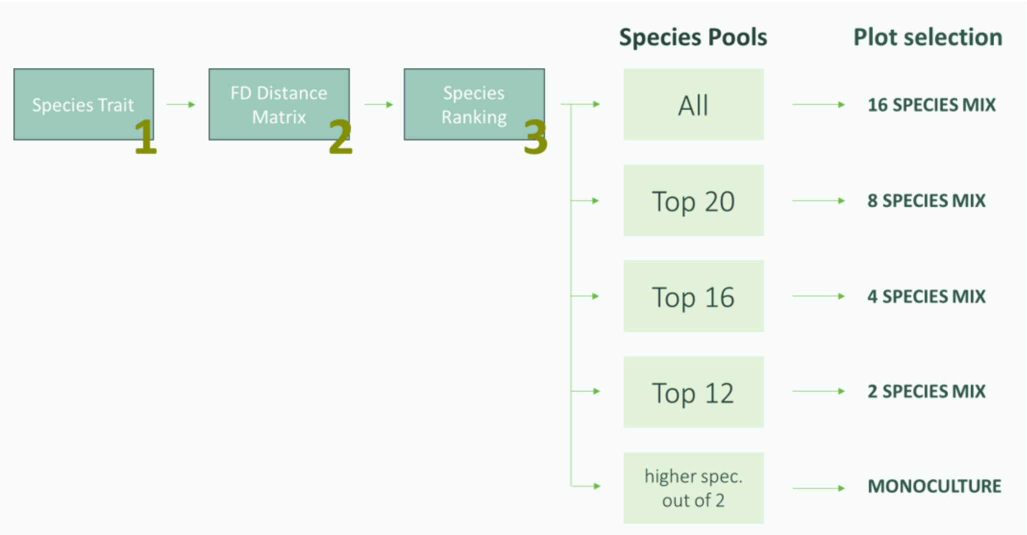


**Figure S 3.** Workflow used to construct the functional-diversity-based extinction pathways.

We started from the species × trait matrix (1) and calculated pairwise functional distances among species (2). Species were then ranked by functional similarity (3), and reduced species pools were defined for each extinction level. For each pool, we retained only those planted BEF-China plots whose species composition was fully contained in the respective pool, resulting in sets of realized 16-, 8-, 4-, and 2-species mixtures and monocultures.
